# Novel Thyroid-Specific Autoantibodies in Patients with Immune-Related Adverse Events Involving the Thyroid Gland

**DOI:** 10.1101/2021.01.14.426670

**Authors:** Ichiro Yamauchi, Akihiro Yasoda, Takuro Hakata, Takafumi Yamashita, Keisho Hirota, Yohei Ueda, Toshihito Fujii, Daisuke Taura, Masakatsu Sone, Nobuya Inagaki

**Author notes:** **Correspondence:** Ichiro Yamauchi, MD, PhD.

## Abstract

**Aims:** Programmed cell death-1 (PD-1) blockade therapy frequently results in immune-related adverse events involving the thyroid gland (thyroid irAEs). Although clinical features of thyroid irAEs are known, the mechanisms remain unclear. Here, we conducted a pilot study to investigate mechanisms of thyroid irAE development from the perspective of autoantibodies.

**Methods:** We performed immunoprecipitation-based assays using sera of 3 patients who developed thyroid irAEs with PD-1 blockade therapy by nivolumab and HEK293T cell lysates, including overexpressed proteins of interest (NKX2-1, PAX8, FOXE1, and HHEX; thyroid-specific transcriptional factors). The pellets were analyzed by western blot to detect the HiBit tag attached to the C-terminus of the proteins.

**Results:** Relevant changes to NKX2-1 bands were not seen in all 3 patients, but PAX8 bands were augmented in patient 2 with lung cancer and patient 3 with renal cell carcinoma. In addition, FOXE1 bands were augmented in patient 1 with malignant melanoma and patient 3, and a HHEX band was augmented in patient 3. Thus, we revealed novel thyroid-specific autoantibodies, PAX8Ab, FOXE1Ab, and HHEXAb. Expression patterns of the antigens recognized by these antibodies were not identical to the primary sites, so autoimmune responses in thyroid irAE may originate from the thyroid gland, and not the malignancy. Considering that TPOAb rather than TgAb is often negative in patients with thyroid irAEs, other mechanisms such as cytotoxic T cell and antigenicity of thyroglobulin may be involved.

**Conclusions:** Although the significance of these novel autoantibodies needs further examination, the present study provides new insights for thyroid autoimmunity.

## Introduction

Immune-related adverse events (irAEs) frequently develop in patients administered immune checkpoint inhibitors. The inhibitors used in clinical practice consist of monoclonal antibodies against cytotoxic T-lymphocyte–associated protein 4 (CTLA-4), programmed cell death-1 (PD-1), and programmed death-ligand 1 (PD-L1). Several endocrine-related organs are involved with irAEs: hypophysitis with CTLA-4 blockade therapy, ACTH deficiency, type 1 diabetes, and thyroid dysfunction with PD-1 blockade therapy (1).

Thyroid irAEs, which are irAEs involving the thyroid gland, are commonly caused by monoclonal antibodies against PD-1 such as nivolumab and pembrolizumab. Our previous case-series study and following retrospective cohort study presented a distinct clinical course of thyroid irAEs: a transient and rapid course of thyrotoxicosis and subsequent persistent hypothyroidism (2,3). High incidence of thyroid irAEs in PD-1 blockade therapy is another important matter: 13.5% in overt thyroid irAEs with nivolumab (3), 14.0–20.8% in thyroid irAEs with pembrolizumab (4–6).

Clinical features of thyroid irAEs are substantially clarified, but mechanisms of thyroid irAE development by PD-1 blockade therapy remain unclear to a large extent. Associations between thyroid diseases and the PD-1/PD-L1 pathway have been reported: both differentiated and anaplastic thyroid cancers express PD-L1, a ligand of the PD-1 receptor (7,8), and a single nucleotide polymorphism (SNP) in the *CD274* gene coding PD-L1 and SNPs in the *PDCD1* gene coding PD-1 associated with Graves’ disease has been identified (9,10). There are no reports regarding associations between SNPs of PD-1/PD-L1 pathway and thyroiditis, a cause of thyroid dysfunction by irAEs (11). We examined and reported that both ligands of PD-1, PD-L1 and PD-L2, are expressed even in normal thyroid tissue (2), which supports the idea that PD-1 blockade therapy could reduce immune tolerance in the thyroid gland.

To investigate further mechanisms of thyroid irAEs, we focused on relationships between thyroid irAEs and prognosis (3,5,12). We previously provided evidence that thyroid irAEs are associated with good prognosis in lung cancer but not in malignant melanoma (3). Here, we wondered antibodies produced by the immune response and their antigens might be intermediators of this prognostic effects. Interestingly, irAEs are likely to develop in the same organs as the primary sites: pneumonitis is common in lung cancer, while skin irAEs are common in malignant melanoma (13). Moreover, patients with pneumonitis as an irAE had longer progression-free survival in non-small cell lung cancer (14), and patients with skin-related irAEs had longer overall survival in malignant melanoma (13). In this context, we hypothesized that antigens common to primary sites and the thyroid gland are involved in the autoimmune responses in thyroid irAEs. In other words, if thyroid irAEs are caused by actions associated with antibodies recognizing antigens common to the primary sites and the thyroid gland, autoimmune responses to cancers could additionally be expected.

Here, to provide mechanistic insights into thyroid irAEs, we conducted a pilot study from the perspective of autoantibodies. The procedure was based on the following findings. Firstly, antibodies related to thyroid irAEs might be different from those of other types of thyroiditis, such as anti-thyroperoxidase antibodies (TPOAbs) and anti-thyroglobulin antibodies (TgAbs), as some patients with thyroid irAEs were double negative for them (3,15,16). Secondly, endocrine-related irAEs including ACTH deficiency and type 1 diabetes as well as thyroid irAEs are commonly associated with severe or null hormonal deficiency. Anti-PIT-1 syndrome suggested us testing transcriptional factors because it presents with severe hormonal deficiency and associates with antibodies for the transcriptional factor specific for cells secreting the hormones (17). In the case of thyroid irAEs, it seemed to be worth examining well-known transcription factors essential for the thyroid gland: NK2 homeobox 1 (NKX2-1), paired box 8 (PAX8), forkhead box E1 (FOXE1), and hematopoietically-expressed homeobox (HHEX) (18). If novel autoantibodies for these thyroid-specific transcriptional factors are detected, the present study provides new insights for exploring the field of thyroid autoimmunity.

## Materials and Methods

### Plasmid construction

We constructed pcDNA3.1 (Thermo Fisher Scientific, Waltham, MA) based plasmid vectors with coding sequences for *NKX2-1* (NM_003317), *PAX8* (NM_003466), *FOXE1* (NM_004473), and *HHEX* (NM_002729) attached to a FLAG tag at the 5’ end and HiBit tag (Promega, Madison, WI) at the 3’ end, using a PCR technique. These coding sequences were derived from the cDNA of human normal thyroid tissue obtained from thyroidectomy specimens for thyroid cancer, as previously described (2). These constructed vectors (pcDNA3.1-FLAG-NKX2-1-HiBit, pcDNA3.1-FLAG-PAX8-HiBit, pcDNA3.1-FLAG-FOXE1-HiBit, and pcDNA3.1-FLAG-HHEX-HiBit) were verified by sequencing.

### Cell culture and protein extraction

HEK293T, a derivative of human embryonic kidney 293 cells, was maintained in Dulbecco’s modified Eagle’s medium (DMEM) (Thermo Fisher Scientific) with Antibiotic Antimycotic (Thermo Fisher Scientific) and 10% fetal bovine serum (Sigma-Aldrich, St. Louis, MO) as previously described (19).

Transient transfections were performed in 100-mm dishes. HEK293T cells were seeded at 1.0 × 10^6^ cells/dish in 10 mL of antibiotic-free DMEM supplemented with 10% fetal bovine serum. After incubation for 24 h, transient transfections of 5 µg of pcDNA3.1-based vectors were performed using 25 µg of PEI MAX (Polysciences, Warrington, PA). After an additional incubation for 24 h, whole cell lysates in radioimmunoprecipitation assay buffer (Nacalai Tesque, Kyoto, Japan) were obtained. We evaluated the protein concentration by the Bradford method with Protein Assay CBB Solution (Nacalai Tesque) using bovine serum albumin as a standard.

### Immunoprecipitation and western blot

We mixed 500 μg of extracted cell lysates and 100 μL of patients’ sera with 100 μL of SureBeads™ Protein G Magnetic Beads (Bio-Rad, Hercules, CA) and added phosphate-buffered saline containing 0.2% of Tween 20 (PBS-T) (Sigma-Aldrich) up to a total of 1000 μL. After rotation for 24 h at room temperature, beads were washed in PBS-T, and bound proteins were eluted with 20 μL of 20 mM Glycine (pH 2.0) (Nacalai Tesque) and neutralized with 2 μL of 0.1 M phosphate buffer (pH 7.4) (Nacalai Tesque).

The total amount of the pellets and 2.5 μg of input proteins were electrophoresed in Bolt 4–12% Bis-Tris Plus Gels (Thermo Fisher Scientific), and transferred onto polyvinylidene difluoride membranes with iBlot Dry Blotting System (Thermo Fisher Scientific), according to the manufacturer’s instructions.

For detecting HiBit-tag, we used the Nano Glo HiBit Blotting System (Promega). After overnight incubation in LgBit protein solution at 4 °C, bands were detected by adding substrate solution in ImageQuant LAS 4000 (GE Healthcare, Chicago, IL). For detecting other tags and proteins, the membranes were blocked with Blocking One (Nacalai Tesque). We incubated with primary antibodies overnight at 4 °C, and then with secondary antibodies for 2 h at room temperature. Bands were detected by a chemiluminescent method with Chemi Lumi One Super (Nacalai Tesque) in ImageQuant LAS 4000. The primary antibodies used were the mouse monoclonal anti FLAG M2 antibody (F3165; Sigma Aldrich) and rabbit polyclonal antibody against β-actin antibody (4967; Cell Signaling Technology, Danvers, MA) as an endogenous control. The secondary antibody used was horseradish-peroxidase (HRP)-conjugated goat polyclonal antibody against mouse IgG1 (1070-05; Southern Biotech, Cambridge, United Kingdom), HRP-conjugated goat polyclonal antibody against rabbit IgG (4050-05; Southern Biotech), and HRP-conjugated goat polyclonal antibody against human Ig (2010-05; Southern Biotech).

## Results

### Case Presentation

In the present study, we examined sera of 3 patients who developed thyroid irAEs. Their detailed characteristics are shown in Table 1 and their clinical courses are described below.

**Table 1.**
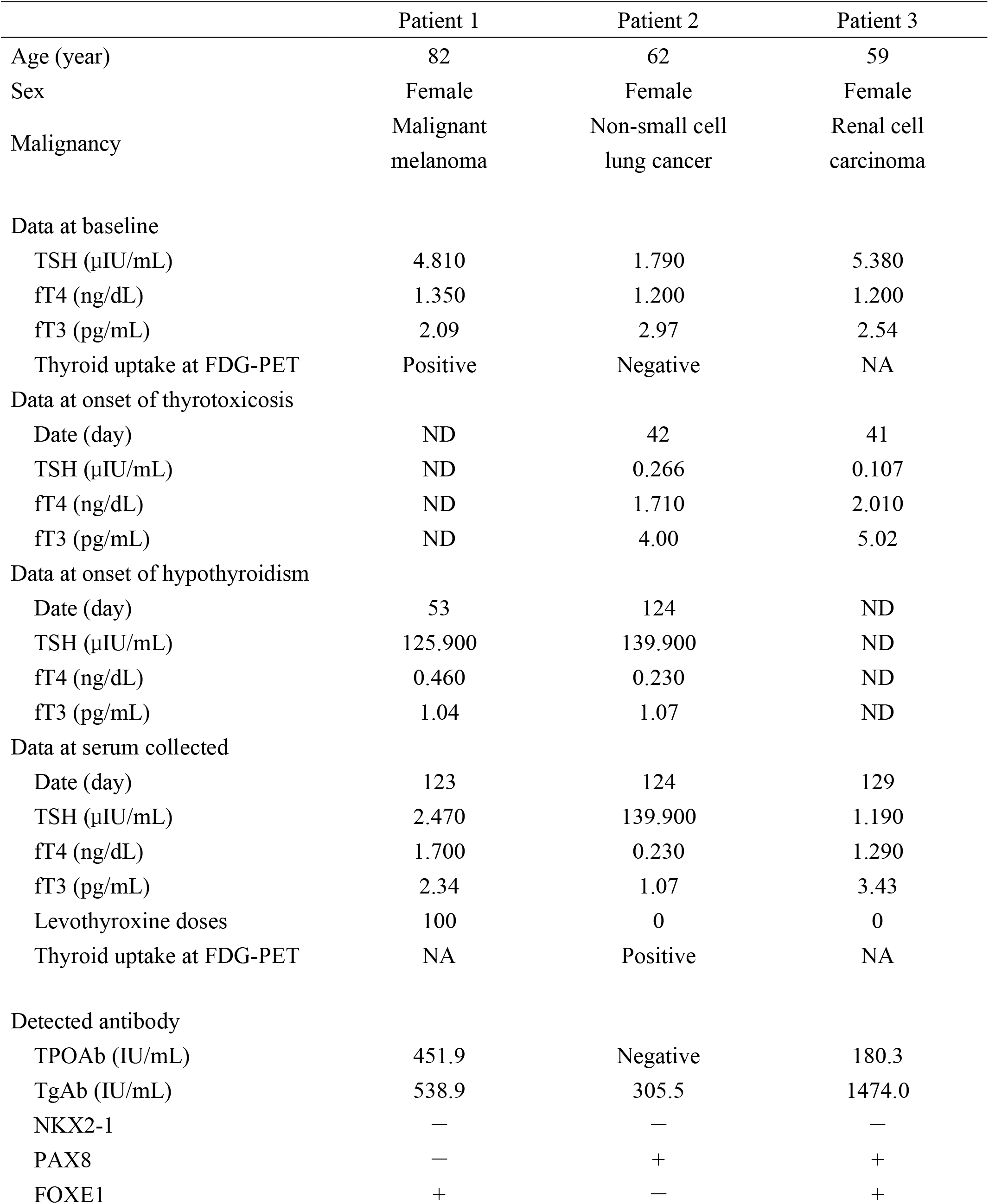
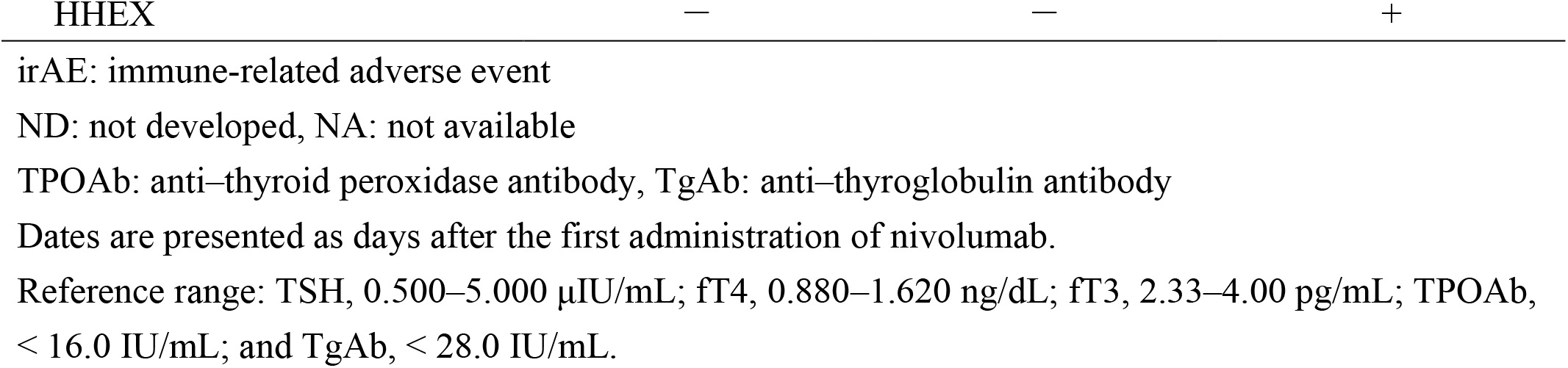
Characteristics of studied cases with thyroid irAEs.

#### Patient 1: 82-year-old woman with malignant melanoma

Four years prior to starting nivolumab therapy, she was diagnosed with primary hypothyroidism and was treated with 50 μg of levothyroxine. Nevertheless, as she continued with 50 μg of levothyroxine therapy, hypothyroidism with fatigue developed 53 days after the first administration of nivolumab. TPOAb and TgAb were double positive at the onset of this hypothyroidism. There was no evidence of thyrotoxicosis during the entire period of nivolumab therapy. We obtained serum samples 123 days after the first administration of nivolumab, even though she was euthyroid at this time due to the increase in levothyroxine dose to 100 μg/day.

#### Patient 2: 62-year-old woman with non-small cell lung cancer

She had no past history of thyroid diseases and was euthyroid before nivolumab therapy. Mild thyrotoxicosis developed 42 days after the first administration of nivolumab. Nivolumab therapy was discontinued because muscle pain and elevation of creatine kinase (1058 IU/L) were observed simultaneously. She then developed asymptomatic hypothyroidism at 124 days after the first administration of nivolumab, and received levothyroxine replacement. TPOAb was negative and TgAb was positive at the onset of hypothyroidism. We obtained serum samples 124 days after the first administration of nivolumab when she developed hypothyroidism.

#### Patient 3: 59-year-old woman with renal cell carcinoma

She had prior axitinib therapy and presented with subclinical hypothyroidism before nivolumab therapy. At 41 days after the first administration of nivolumab, she developed thyrotoxicosis that worsened and caused fatigue. TPOAb and TgAb were double positive at the onset of thyrotoxicosis. Prolonged diarrhea and elevated liver enzymes (AST 366 IU/L, ALT 329 IU/L, Grade 3) were also observed. For treatment of these irAEs, glucocorticoids were administered; 100 mg/day methylprednisolone for 4 days, 50 mg/day prednisolone for 3 days, and 25 mg/day prednisolone for 3 days. Her condition, liver enzymes, and abnormalities of thyroid function improved without developing hypothyroidism. We obtained serum samples 129 days after the first administration of nivolumab, even though she was euthyroid at this time.

### Identification of autoantibodies in patients’ sera

We initially confirmed that plasmid vector constructs successfully expressed proteins of interest (Figure 1). Detection of FLAG-tag attached to the N-terminus and HiBit tag to the C-terminus showed similar results that each band corresponded to its putative molecular weight; the band of FLAG-NKX2-1-HiBit was at 45 kDa, the bands of FLAG-PAX8-HiBit bands were at 52 and 55 kDa, the FOXE1 band was at 44 kDa, and the HHEX band was at 42 kDa.

**Figure 1.**
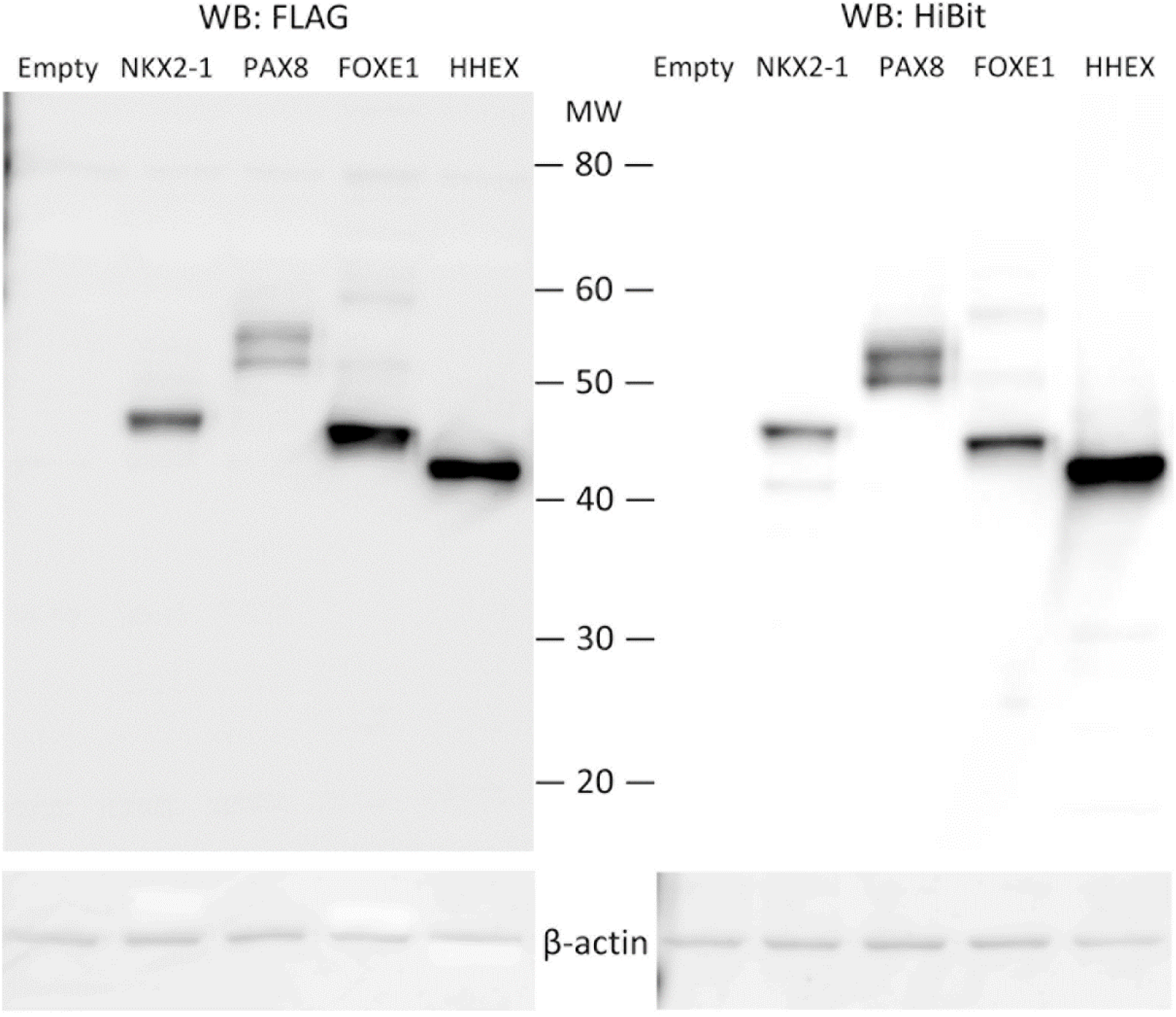
Verification of constructed plasmid vectors. Protein lysates of transfected HEK293T cells were analyzed by western blot. Transfected plasmid vectors are presented as follows: Empty, pcDNA3.1; NKX2-1, pcDNA3.1-FLAG-NKX2-1-HiBit; PAX8, pcDNA3.1-FLAG-PAX8-HiBit; FOXE1, pcDNA3.1-FLAG-FOXE1-HiBit; and HHEX, pcDNA3.1-FLAG-HHEX-HiBit. The left panel shows detection of the FLAG tag attached to the N-terminus, and the right panel shows detection of the HiBit tag attached to the C-terminus. Molecular weights (MW) ranged from 20 to 80 kDa.

Subsequently, we performed immunoprecipitation using cell lysates including overexpressed proteins of interest and sera of patients (Figure 2). We prepared control pellets obtained by using serum of an individual control subject who was confirmed as euthyroid and negative for TPOAb and TgAb, defined as control A. Immunoprecipitated pellets were analyzed by western blot. There were no obvious differences in detected human immunoglobulin (Ig) among the pellets of the three patients and the control A.

**Figure 2.**
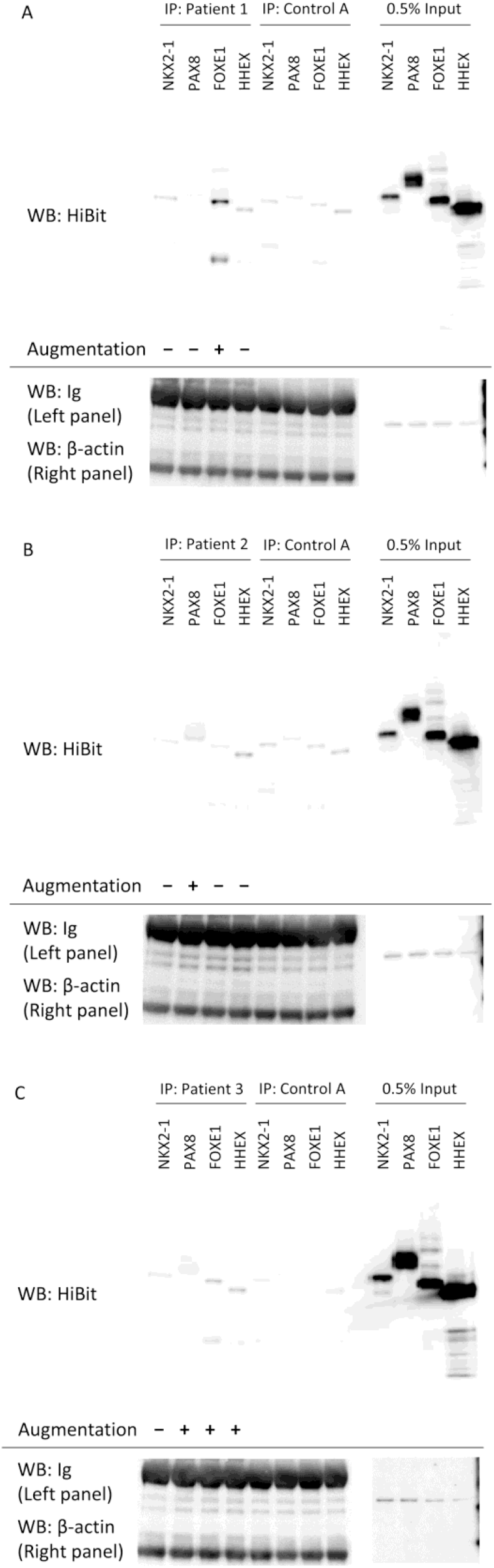
Western blot of pellets of immunoprecipitation (IP) using cell lysates overexpressing proteins of interest and sera of patients. Tagged proteins of NKX2-1, PAX8, FOXE1, and HHEX were individually overexpressed. A pellet of control A was obtained using serum of a subject that was confirmed as euthyroid and negative for TPOAb and TgAb. We electrophoresed 0.5% of input protein as size markers of the overexpressed protein, and detected human immunoglobulins (Ig) and β-actin as controls of IP and protein loading, respectively. **(A)** Patient 1 with malignant melanoma. Compared to the control A, the FOXE1 bands were augmented. **(B)** Patient 2 with non-small cell lung cancer. The PAX8 band was augmented. **(C)** Patient 3 with renal cell carcinoma. The PAX8 band, FOXE1 bands, and HHEX band were augmented.

We evaluated augmentation of bands, suggesting that antibodies were bound to the corresponding proteins. The HiBit tag was used because of its low background signals. Augmentation of the NKX2-1 bands were not seen. The lower bands of PAX8 were augmented in patients 2 and 3 (Figure 2B and Figure 2C, respectively), while the upper bands showed no changes. For FOXE1, bands at 44 kDa and additional bands at 25 kDa (detected only in pellets) were augmented in patients 1 and 3 (Figure 2A and Figure 2C, respectively). A HHEX band was augmented in patient 3 (Figure 2C). These results are summarized in Table 1.

Finally, to reinforce specificity of augmentation of PAX8, FOXE1, and HHEX bands in these 3 patients, we additionally tested sera of 3 subjects who was confirmed as euthyroid and negative for TPOAb and TgAb, defined as control B, C, and D. As expected, no obvious augmentation of bands compared to control A was observed in these additional controls (Figure 3).

**Figure 3.**
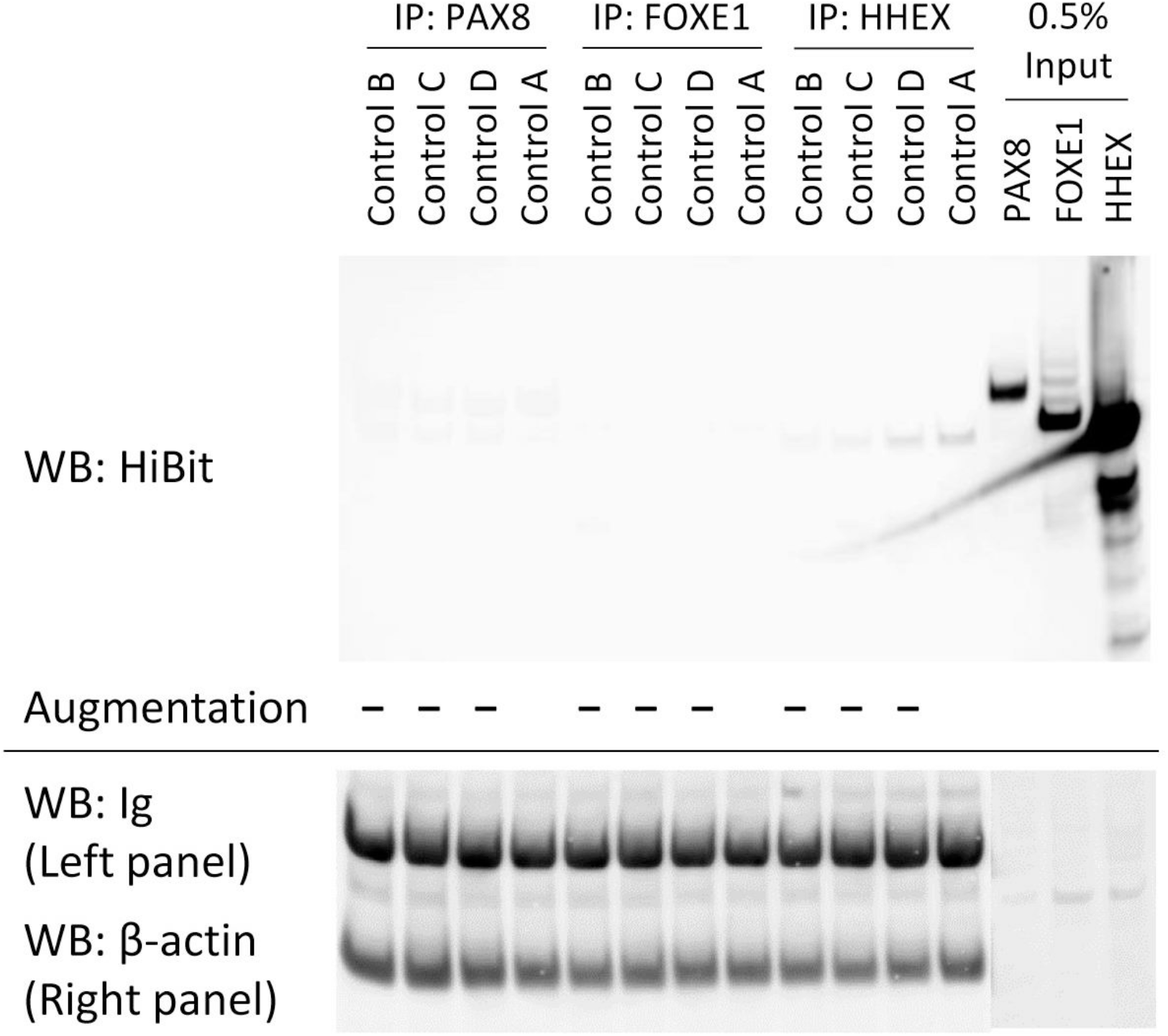
Western blot of IP pellets using cell lysates overexpressing proteins of interest and sera of 3 additional control subjects referred to as control B, C, and D. Compared to the control A identical to that in Figure 2, the PAX8 band, FOXE1 bands, and HHEX band were not augmented in any samples.

## Discussion

Here we performed a pilot study to investigate mechanisms of thyroid irAEs from the perspective of autoantibodies. Immunoprecipitation-based assays using sera from 3 patients who developed thyroid irAEs revealed novel autoantibodies for PAX8, FOXE1, and HHEX, which are thyroid-specific transcriptional factors.

Reliability of our experimental results was supported by the following facts. Augmentation of bands observed differently; this would be in contrast to systemic activated immunoreactivity if all four bands were augmented. In addition, we verified that the amounts of immunoglobulin (Ig) collected in IP pellets did not differ between patients and a control; if the Ig amount had been different, this would indicate non-specific immunoreactivity. Lastly, 3 additional controls did not present augmentation of bands; novel autoantibodies seemed to be uncommon at least in normal subjects.

There are some autoantibodies involving thyroiditis, but thyroid-specific autoantibodies are limited to TPOAb and TgAb. TPOAb and TgAb are mainly seen in chronic autoimmune thyroiditis (Hashimoto’s thyroiditis). The positive rate of TPOAb and TgAb tests in chronic autoimmune thyroiditis is difficult to know because its diagnosis depends on positive results of TPOAb and/or TgAb. However, we often see patients with hypothyroidism who are double negative for TPOAb and TgAb. A previous study has shown that TPOAb and/or TgAb are positive in more than 90% of patients with chronic autoimmune thyroiditis (20). In other words, there are particular patients without TPOAb and TgAb with autoimmune thyroiditis.

Interestingly, patients without TPOAb and TgAb at the development of thyroid irAEs are frequently observed (3,15,16). For example, of 17 patients with thyroid irAEs in our previous study, 7 patients were double positive, 5 were double negative, and 5 were exclusively positive for TgAbs, while no patients were exclusively positive for TPOAbs (3). Even at baseline, similar discrepancy that positivity of TgAb was higher than that of TPOAb was seen in patients with thyroid irAEs (21). The etiology may differ between chronic autoimmune thyroiditis and thyroid irAE by PD-1 blockade therapy.

The present investigation focused on thyroid-specific transcriptional factors by reasons discussed in the Introduction. Autoantibodies for transcriptional factors were reported in some papers such as PIT1-antibody in hypopituitarism (17) and TIF1-γ in dermatomyositis (22). In addition, antibodies recognizing intracellular antigens such as Hu, Yo, and Ri are often detected in paraneoplastic syndrome (23). However, administration of these antibodies to animals did not show relevant pathogenicity (24–26). On the other hand, the involvement of cytotoxic T cells positive for CD8 was suggested in paraneoplastic syndrome related to the Hu antibody (27). CD8 T cells were increased in peripheral blood after PD-1-targeted therapy in lung cancer patients (28). The cytotoxicity of T cells might have a more dominant role than antibody-specific immunoglobulins in irAEs by PD-1 blockade therapy.

To discuss the significance of the novel autoantibodies, their origins are an important matter. PAX8 was not expressed in malignant melanoma, rarely in lung cancer, but often in renal cell carcinoma (29). FOXE1 was often expressed in lung cancer (30). However, our results did not correspond with these expression patterns: PAX8 antibodies were found in patient 2 with lung cancer and in patient 3 with renal cell carcinoma, and FOXE1 antibodies were found in patient 1 with malignant melanoma and patient 3 with renal cell carcinoma. It is well known that the adult thyroid gland expresses PAX8 and FOXE1 (29,31). In this context, autoimmune responses in thyroid irAEs might originate from the thyroid gland.

We believe that the thyroid gland is susceptible to an autoimmune response. This is supported by the high frequency of chronic autoimmune thyroiditis (32). This cause has not been completely understood, but the following information seems to be suggestive of it. Thyroglobulin in the thyroid follicle may have high antigenicity because thyroglobulin-immunized mice often develop thyroiditis even if the mice were immunized by murine thyroglobulin (33). The thyroid gland needs strong immune tolerance because the thyroid gland expresses both PD-L1 and PD-L2 (2). Moreover, TgAb rather than TPOAb could be associated with thyroid irAEs by PD-1 blockade therapy (3,21), including through the prediction of thyroid irAEs.

Integrating the findings so far, we consider putative mechanisms of thyroid irAEs: 1) positive TgAb may reflect preexisting autoimmune response because thyroglobulin has potential for antigenicity; 2) PD-1 blockade therapy easily causes thyroiditis by disrupting the upregulated immune tolerance in the thyroid gland; 3) this type of thyroiditis is accompanied by the production of thyroid-specific autoantibodies except for TPOAb and TgAb, but the pathogenicity does not depend on these autoantibodies; 4) in terms of a relationship between thyroid irAEs and prognosis, immune response by CD8 T cells occurs if tumors express antigens recognized by thyroid-specific autoantibodies.

Several unresolved issues remain after this pilot study. Testing larger number of samples including various controls such as chronic autoimmune thyroiditis occurred without immune checkpoint therapy is essential for confirming specificity of novel autoantibodies. Whether patients without TPOAbs and TgAbs at the development of thyroid irAEs have these novel antibodies is an important matter, because our tested patients had TPOAbs and/or TgAbs. As for assays, it is better to improve background signals observed in the controls and low throughput. Although enzyme immunoassay might be suitable for this aim, specificity of the signals should be validated by western blot analysis while referring to the present study. Similar examinations for other thyroid-specific proteins may also contribute to clarify mechanisms of thyroid irAE.

In this pilot study, we identified novel thyroid-specific autoantibodies, PAX8Ab, FOXE1Ab, and HHEXAb in some patients with thyroid irAEs by PD-1 blockade therapy. Although the significance related to pathogenicity and specificity requires further validation, our finding provides insights for investigators of thyroid immunity.

## Data Availability Statement

All datasets presented in this study are included in the article/ supplementary material.

## Ethics Statement

The studies involving human participants were reviewed and approved by the Institutional Review Board and Ethics Committee of the Kyoto University Graduate School of Medicine. Written informed consent for participation was not required for this study in accordance with the national legislation and the institutional requirements.

## Author Contributions

IY designed and conducted experiments. AY, TH, TY, KH, YU, TF, DT, and MS contributed to the discussion. AY and NI provided funding and supervised the entire study. IY drafted the manuscript, and all authors reviewed, edited, and approved the manuscript.

